# Neutrophil extracellular traps impair regeneration

**DOI:** 10.1101/2020.07.06.189910

**Authors:** Eric Wier, Mayumi Asada, Gaofeng Wang, Martin P. Alphonse, Ang Li, Chase Hintelmann, Christine Youn, Brittany Pielstick, Roger Ortines, Lloyd S. Miller, Nathan K. Archer, Luis A. Garza

## Abstract

Fibrosis is a major health burden across diseases and organs. To remedy this, we study wound induced hair follicle regeneration (WIHN) as a model of non-fibrotic healing that recapitulates embryogenesis for de novo hair follicle morphogenesis after wounding. We have previously demonstrated that TLR3 promotes WIHN through binding dsRNA, but the source of which is still unclear. Here, we demonstrate that multiple distinct contexts of high WIHN all show a strong neutrophil signature, and given the likelihood of nuclear dsRNA release during the production of neutrophil extracellular trap (NETs), we hypothesized that mature neutrophils and NETs might promote WIHN. Consistent with this, in addition to the presence of mature neutrophils shortly after wounding, neutrophils remain within the wound after the barrier is reestablished, where they produce extracellular traps (NETs) that likely release spliceosomal U1 dsRNA. Contrary to our hypothesis, genetic models of neutrophil depletion show enhanced WIHN. Pad4 null mice that are defective in NET production also augment WIHN. Finally, using single-cell RNA sequencing, we identified a dramatic increase in mature neutrophils in the wound beds of low regenerating Tlr3-/- mice. Taken together, these results demonstrate that although mature neutrophils are stimulated by a common pro-regenerative cue, their presence and NETs hinder WIHN.

## Introduction

After suffering a wound, the body initiates a well-coordinated physiological process to restore homeostasis and reestablish the barrier. This spontaneous process comprises four discrete phases: hemostasis, inflammation, proliferation, and remodeling^1^. Although all of these phases have been extensively studied, the molecular details as to why repair tends to result in fibrotic scar tissue rather than complete regeneration have not been fully elucidated. This tendency of wound repair to lead to fibrosis, and in some cases, hypertrophic scars, contributes an enormous burden on human health^1-3^. Inflammation and different components of the immune system have been shown to contribute to regeneration in salamanders and zebrafish, as well as to promote healing and maintain barrier function in mammals upon mucosal injury^4-6^. Although the cellular effects of macrophages and T cells are well studied^7-10^, the role of neutrophils in modulating regeneration remains elusive.

Unlike urodele salamanders, complete skin regeneration after wounding in mammals is rare^11^. It occurs via de novo hair follicle generation in mice and rabbits through a process that mimics skin embryogenesis^12-15^. This process of de novo follicle neogenesis (Wound Induced Hair Neogenesis; WIHN) was first fully characterized in mice after receiving full thickness wounds^16^. These regenerated follicles establish a distinct stem cell population, express characteristic differentiation markers, produce functional hair shafts, and can complete the hair cycle. In addition to the hair follicles, sebaceous glands, specialized vascular and nerve supports, and surrounding fat cells are regenerated^16,17^.

Immediately after wounding, a robust inflammatory phase occurs, which allows the ingress of keratinocytes and fibroblasts to proceed afterward. The early stage of wound healing is defined by the dramatic recruitment of mature neutrophils, which are instrumental in providing defense against microbial pathogens^18-22^. This is followed by an influx of macrophages (Mϕ’s) that continue the phagocytic processes begun by neutrophils and aid in the transition to the proliferative phase of wound healing^23-25^. While much is known about the importance of this inflammatory phase for preventing and abrogating infections, less is known about how it influences regenerative capacity or WIHN. Increasingly, macrophages have been shown to be essential for WIHN via TNF-induced AKT/ β-catenin signaling^8-10^. However, the role of neutrophils in modeling regeneration or WIHN is unclear.

Toll-like receptors (TLRs) are highly conserved single-pass membrane-spanning receptors that recognize structurally conserved molecular components of invading microbes and activate a cascade of inflammatory signaling pathways^26^. Rather than simply recognizing pathogen associated molecules, they can also initiate “sterile” inflammation upon recognizing damage-associated molecular patterns (DAMPs), which are critical to recruit immune cells and initiate wound healing^27^. TLR3 is activated by dsRNA and has primarily been studied in the context of viral infection^28^. Mounting evidence shows that TLR3 also plays an important role in wound repair^29-37^. Synthetic double strand RNA (dsRNA) polyriboinosinic-polyribocytidylic acid (poly(I:C)) treatment dramatically increases WIHN in mice. Furthermore, wound-released dsRNA activates TLR3 to promote hair follicle regeneration^34^. The identity and source of the physiological RNA remains an open and important question in the field. Notably, the dsRNA U1 spliceosomal small nuclear RNA (snRNA), may be an important endogenous RNA sensed via TLR3^32,33,38,39^. Specifically, UV damage releases U1 snRNA that stimulates cytokine production in keratinocytes and increases barrier gene transcription^32,33^.

In addition to phagocytosis and degranulation, mature neutrophils can produce extracellular traps (ET), large extracellular web-like structures composed of decondensed chromatin bound to various cytosolic and granule proteins^21,22,40-42^. While originally recognized as a defense mechanism against pathogens^21,40,41^, they have also been found to mediate sterile inflammatory processes^43,44^. In the absence of infection, ETs can be stimulated in sterile tissue environments through various cytokines^41,45-47^ and by activated platelets^43,44^. Interestingly, ETs are found within sterile wounds of mice and delay wound healing^48^. Mechanistically, ETs are formed by the rapid decondensation of the cellular chromatin, followed by the fragmentation of the nuclear membrane and mixing of the nuclear and cytoplasmic compartments, before being expelled from the cell. The ability of neutrophils to rapidly migrate to the wound site and produce ETs, coupled with the nuclear localization of some dsRNA, made us question whether neutrophils were a source of the dsRNA critical for WIHN. Interestingly, while there are extensive studies on the DNA components released during ET formation^22,42,49^, the RNA components are poorly understood.

To probe how mature neutrophils influence wound regeneration and WIHN, we analyzed multiple microarrays from distinct contexts of high regenerating mice and found a common neutrophil signature. Using immunofluorescence and flow cytometry we found that neutrophils remain in the wound bed, albeit at low levels, after the acute inflammatory phase, where they produce NETs that contain the nuclear U1 dsRNA. To define how this influences regeneration, we used antibody mediated neutrophil depletion to eliminate neutrophils from the wound bed at select time points, but discovered that the technique is ineffective in the context of large full-thickness wounds, which are necessary to initiate WIHN. However, using a neutrophil-specific diphtheria toxin ablation model, we were able to deplete mature neutrophils in the wound bed and found that—contrary to our initial hypothesis--the absence of mature neutrophils enhances WIHN. Eliminating neutrophil’s ability to produce NETs by knocking out Pad4 also boosted WIHN, confirming the negative influence of neutrophils on regeneration. Finally, we used single-cell RNA sequencing to characterized WIHN deficient Tlr3-/- mice and found that they have a dramatically increased population of mature neutrophils in the re-epithelized wound bed, compared to wild-type mice, likely contributing to their diminished regenerative capacity. These results indicate that, while important for preventing infection, mature neutrophils and their NETs negatively impact regeneration and WIHN. Although a common pro-regenerative signal might increase neutrophil infiltration, mature neutrophils instead likely contribute to fibrosis.

## Results

### Neutrophil signature present during skin regeneration

To begin to characterize the role of neutrophils in WIHN, we performed bioinformatic analysis on previous microarrays of multiple distinct high regenerating mouse models, probing for innate immune and neutrophil signatures. First, we analyzed the proteome comparing the center of the re-epithelialized wound bed (high WIHN) to the surrounding periphery/edge (low WIHN) (Fig. 1a)^37^. Gene ontology analysis revealed that, in the area of high WIHN, neutrophil aggregation and other defense pathways against bacterium are enriched, characterized by an abundance of antimicrobial and granular proteins, such as Neutrophil Elastase (Elane), Cathelicidin (Camp), and Myeloperoxidase (Mpo) (Fig. 1b-c). In a second model system, high WIHN Rnasel-/- mice (manuscript in review), at the time of scab detachment, are also enriched in genes associated with neutrophils. Neutrophil chemotaxis is the most significant upregulated gene ontology category, with other chemotactic and inflammatory pathways up as well (Fig. 1d). Finally, we analyzed the gene expression changes between wounded specific pathogen free (SPF) mice, which have increased regeneration and WIHN, when compared to germ free (GF) mice (manuscript in review), at the time of scab detachment. Like the other two high regeneration models, when compared to GF, SPF mice had elevated neutrophil chemotaxis and immune response transcripts (Fig. 1e). Together, these disparate experimental contexts demonstrate that neutrophil chemotaxis can correlate with high regeneration and WIHN.

**Figure 1:**
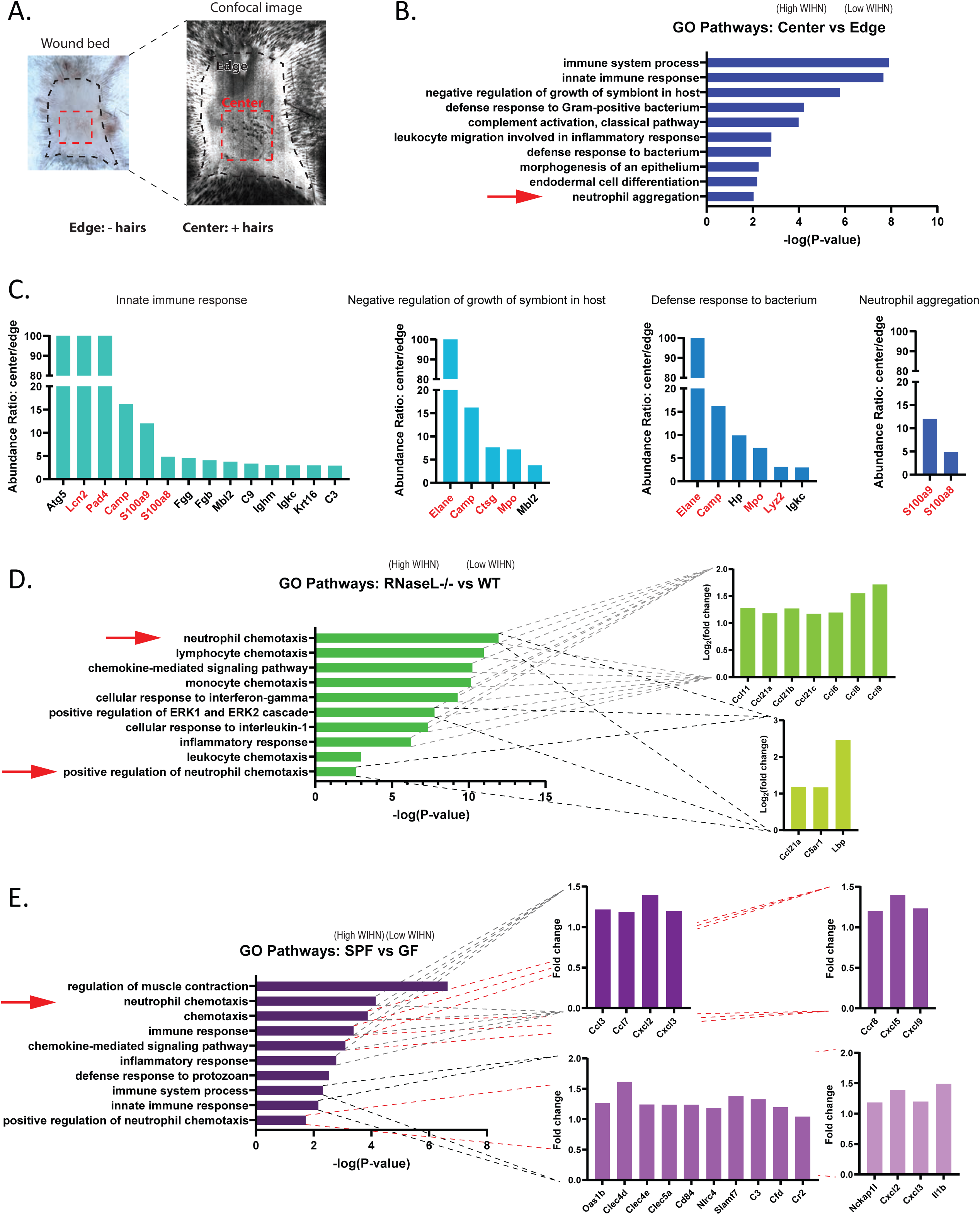
Neutrophil signature correlates with high skin regeneration in multiple models. **a**. Schematic of hair neogenesis preferential localization to wound center (high WIHN) rather than wound edge (low WIHN). **b**. Proteomic gene ontology (GO) analysis of the top 100 genes wound center versus wound edge in wild type mice (at scab detachment) shows a predominance in innate immune response pathways and neutrophil signatures. **c**. Abundance ratios of genes from select GO terms highlighted in b. show enrichment of antimicrobial and granular proteins, labeled in red. **d**. GO analysis of the top 200 genes from high WIHN Rnasel^-/-^ versus wild-type mice at scab detachment shows a predominance of neutrophil and innate immune cell chemotaxis pathways. Inset graphs show the gene fold expression changes for genes present in that category. **e**. Neutrophil chemotaxis and innate immune categories are included in Gene Ontology (GO) enrichment analysis of the top versus bottom 500 differentially expressed genes between Specific pathogen-free (SPF; high WIHN) and germ-free (GF; low WIHN) mice on ∼WD12 (scab detachment) wound beds. (n=3 independent animals per group). Inset graphs show the gene fold expression changes for genes present in that category.

### Neutrophils persist in the wound bed after the acute inflammatory phase, producing extracellular traps

Given the above correlations of high neutrophil infiltration to high WIHN, we first characterized mature neutrophil infiltration in the wound beds of C57BL/6J mice after a large square wound cut into the center of their backs, to the depth of fascia. As anticipated, neutrophils are abundant in the acute phase of the wound healing process (1-3 days post wounding), as seen by H&E staining (Fig. 2a) and immunofluorescence of Myeloperoxidase (MPO), a major neutrophil granule protein (Fig. 2b). While neutrophils predominate at early time points, they are still present in the wound bed as late as wound day 11, well after re-epithelization has completed (Fig. 2c). In contrast, macrophages only accumulate starting at wound day 3 (Fig. 2b) but remain a major component of the wound even after re-epithelization (Fig. S1). These results show the dynamic changes of immune cell infiltration, but with an underappreciated persistence of some neutrophils late in wound healing, during the time of morphogenesis.

**Figure 2:**
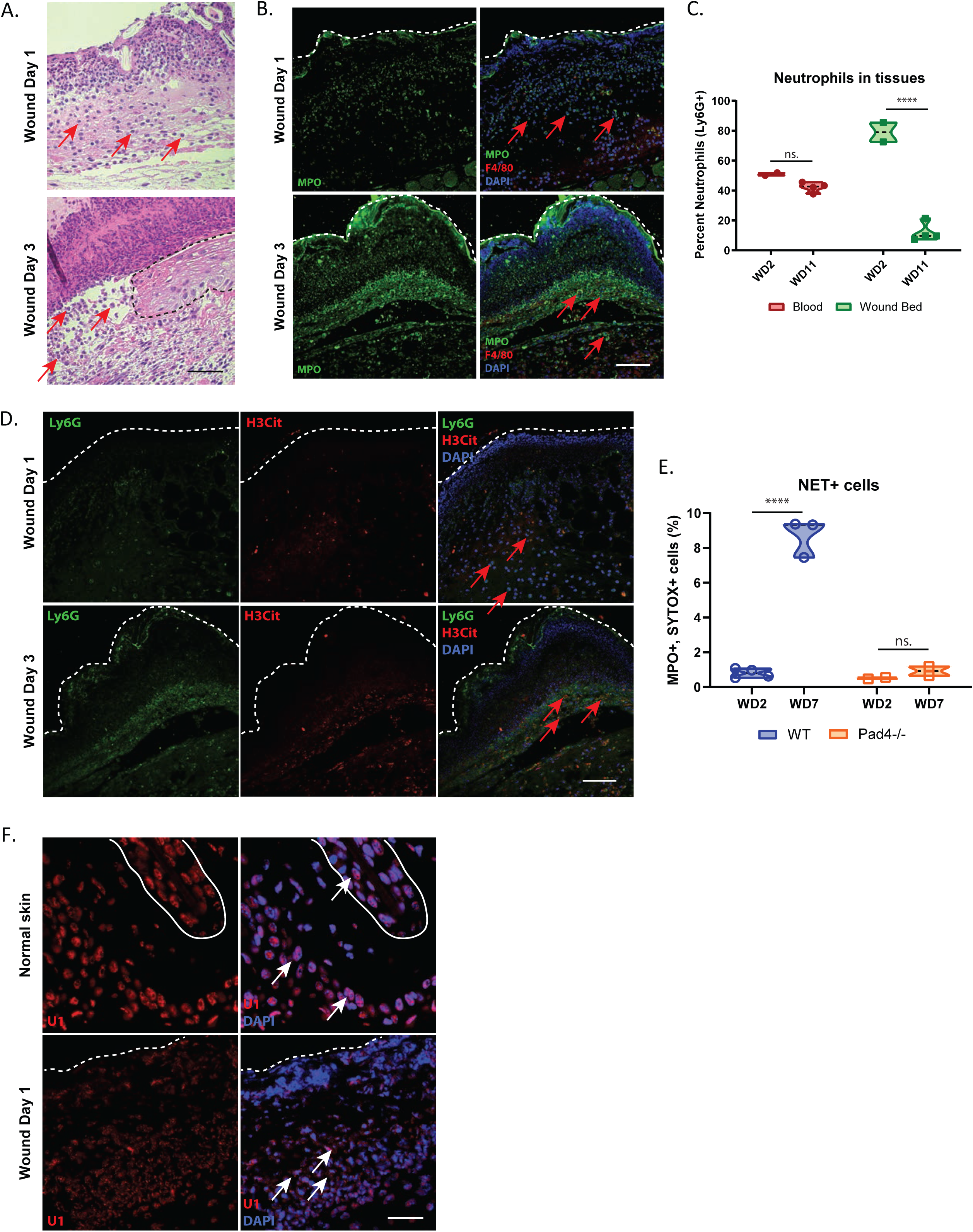
Neutrophils persist in wound bed after the acute inflammatory phase, producing extracellular traps. **a**. Neutrophils are present in the wound beds of C57BL/6J mice at early time points, visible in representative hematoxylin and eosin [H&E] staining. Red arrows show select neutrophils, evident by their multilobular nuclei. The black dashed line signifies the migrating epithelial front. Black scale bar = 50 µm. **b**. Neutrophils predominate throughout the wound beds of C57BL/6J mice on wound days 1 and 3, visible in prominent MPO immunofluorescence (green). Few macrophages are present (Red, F4/80). Red arrows show select neutrophils. The white dashed line signifies the dorsal edge of the wound bed. White scale bar = 200 µm. **c**. Percent neutrophil (Ly6G+ cells from total CD45+ cells) levels are consistent in the blood throughout the wound time course, but drop in the wound bed at wound day 11, as measured by FACS. ****p < 0.0001, as calculated by two-way ANOVA. n.s., not significant. N = 2 vs 4. Results are representative of at least two independent experiments. **d**. Citrullinated histone H3 (H3Cit, red) co-localized with Ly6G+ neutrophils (green), beginning at wound day 3 in the wound beds of immunofluorescence stained C57BL/6J mice, indicating the formation of extracellular traps. Red arrows show select neutrophils. The white dashed line signifies the dorsal edge of the wound bed. White scale bar = 200 µm. **e**. Neutrophil extracellular trap positive cells (MPO+, SYTOX green +) are present at late wound time points, but are absent in the wound beds of PAD4-/- mice, as measured by FACs ****p < 0.0001, as calculated by two-way ANOVA. n.s., not significant. N = 7 vs 4. Results are representative of at least two independent experiments. **f**.Cytoplasmic U1 snRNA is present in the wound bed of C57BL/6J mice, while it localized exclusively in the nuclei of unwounded controls, as visualized by representative FISH. White arrows show select cells with nuclear or cytoplasmic U1 snRNA. The white dashed line signifies the dorsal edge of the wound bed. The solid white line delineates a hair follicle. White scale bar = 80 µm

Given the importance of dsRNA to promote WIHN, and the association of neutrophils with high WIHN, we hypothesized that neutrophil extracellular traps (NETs) release of neutrophil nuclear content might release dsRNA, in addition to dsDNA, from the nucleus to promote WIHN. We visualized citrullinated histone H3 (H3Cit) to identify NETS as early as wound day 3 (Fig. 2d). The production of NETs is mediated by Pad4, an enzyme that modifies the arginine residues on histones to citrulline, which changes their charge, leading to massive chromatin decondensation^50,51^. Consistent with this, NETs are virtually absent in mice lacking Pad4, while abundant in the wound bed after the acute phase of wound healing in WT mice (Fig. 2e). Although NETs are characterized by their extruded DNA, which forms a web-like scaffold containing cytosolic and granular proteins, little is known about the RNA content within them and how that might influence wound healing. Given that U1 small nuclear (sn) RNA is proposed as a TLR3 agonist damage-associated molecular pattern important for skin barrier repair^32,33^, we visualized it by FISH. U1 snRNA is present throughout the wound beds of C57BL/6J mice (Fig. 2f). Rather than the baseline nuclear appearance of unwounded tissue, U1 snRNA in the wound bed is markedly cytoplasmic, suggesting that its cellular localizations shifts from the nucleus as a potential step towards cellular release. Consistent with this, we noted a fine haze of signal extracellularly. Taken together, these data suggest a potential model where neutrophils persist to the morphogenesis stage of wound healing and release extracellular traps that contain U1 dsRNA to perhaps modulate WIHN.

### Large full-thickness wounds dramatically reduce antibody-mediated neutrophil depletion

To functionally assess how neutrophils modulate wound induced hair neogenesis (WIHN), we sought to deplete neutrophils throughout the wound bed at select times. To accomplish this, we adapted a widely used antibody mediated depletion scheme, where we injected neutrophil specific antibodies intraperitoneally (Fig. 3a)^52^. Most experiments were completed using a Ly6G positive monoclonal antibody (1A8 clone), which recognizes a 21-25kD glycosylphosphatidylinositol (GPI)-linked differentiation antigen that is expressed by myeloid-derived cells. While monocytes transiently express Ly6G during bone marrow development, Ly6G expression in peripheral neutrophils directly correlates its level of differentiation and maturation. The less specific anti-granulocyte receptor-1 (Gr-1) antibody (RB6-8C5 clone), which recognizes both Ly6G and Ly6C and thereby also targets dendritic cells and subpopulations of lymphocytes and monocytes, was also used in select experiments. To prevent confounding flow cytometry measurements caused by epitope masking from the initial Ly6G antibody injected during the antibody mediated neutrophil depletion experiments^53^, a distinct Ly6G antibody clone was used for analysis via flow cytometry. Injections were done one day before and one day after large full-thickness wounds on the back of mice. The depletion scheme focused on the early time points of the wound healing process because those were shown to be most critical in dsRNA-Tlr3 enhanced WIHN^34^.

**Figure 3:**
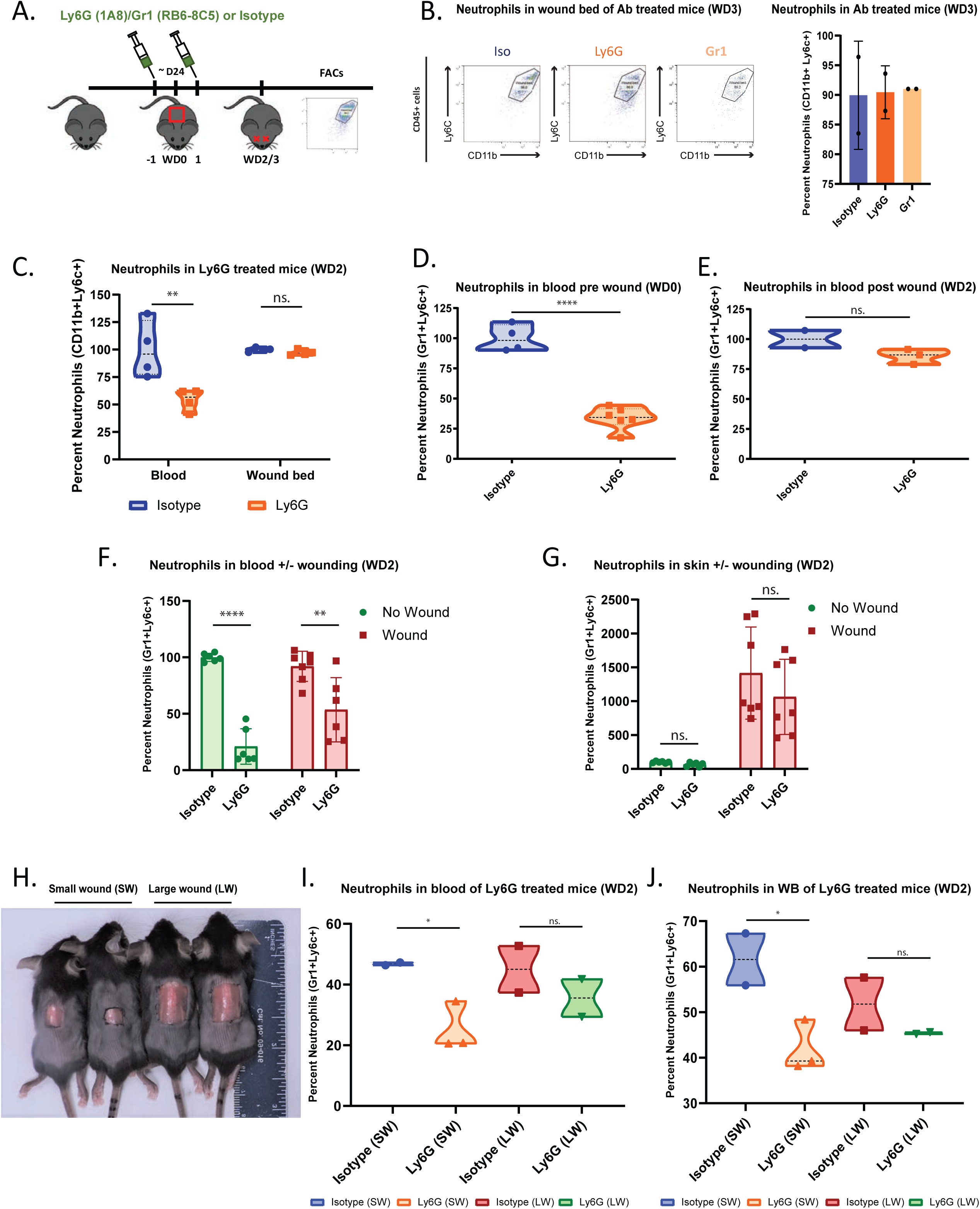
Large full-thickness wounds dramatically reduce antibody-mediated neutrophil depletion. **a**. Schematic of neutrophil depletion via intraperitoneal injection (IP) of Ly6G antibody (1A8 clone, 500μg), Gr1 (Ly6G/Ly6C, RB6-8C5 clone, 200ng), or isotype control. IP injections are done one day before and one day after wounding C57BL/J6 mice with 1.25cm^2^ square wounds. Mice were then sacrificed wound day 2 or 2 for FACs analysis of blood or wound bed (WB). **b**. Antibody mediated depletion is not successful in the wound bed. Representative flow plots from the wound beds (wound day 3) of mice treated as in **a**. The percent neutrophils (CD11b+, Ly6c+) present in these samples are presented to the right. The difference between samples is statistically insignificant, as calculated by the two-tailed Student’s t-test. N = 2. Results are representative of at least two independent experiments. **c**. Antibody mediated. Neutrophil depletion is marginally successful in the blood, but not the wound bed, two days after wounding. The percent neutrophils (CD11b+, Ly6c+), at wound day 2, present in the blood and wound beds of mice treated as in **a**. (Ly6G, 1A8). **p < 0.0011, as calculated by two-way ANOVA. n.s., not significant. N = 4. Results are representative of at least two independent experiments. **d**. Neutrophil depletion from the blood is successful 1-day post injection (Ly6G, 1A8), in the absence of wounding, via FACs. The neutrophil depletion is statistically significant. ****p < 0.0001, as calculated by two-tailed Student’s t-test. N = 4 vs 6. **e**. The efficacy of neutrophil depletion from the blood diminishes after 1.25×1.25cm^2^ wounds are inflicted. The percent neutrophils (Gr1+, Ly6c+) in the blood at wound day 2, via FACs. The neutrophil depletion is statistically insignificant. n.s., not significant. N = 2 vs 3. Results are representative of at least three independent experiments. **f**. Neutrophil depletion from the blood is more pronounced in unwounded mice. The percent neutrophils (Gr1+, Ly6c+) present in the blood at wound day 2 of mice IP injected with Ly6G antibody (1A8, 500 μg), with or without wounding, via FACs. ****p < 0.0001 vs **p=0.0038 as calculated by two-way ANOVA. N = 6 vs 7. **g**. Neutrophil depletion from the skin is ineffective, regardless of wound status. The percent neutrophils (Gr1+, Ly6c+) present in the wound bed or skin at wound day 2 of mice IP injected with Ly6G antibody (1A8, 500ug), with or without wounding. The neutrophil depletion is insignificant in both cases, as calculated by two-way ANOVA. n.s., not significant. N = 6 vs 7. **h**. C57BL/6J mice were given full thickness wounds on the center of their backs. The normal large wound (LW) is 1.25×1.25cm^2^, while the smaller wound (SW) is 0.6×0.6cm^2^. **i**. Neutrophil depletion from the blood is successful on small, but not large wounded mice. Mice wounded with small or large wounds (**h**.) were IP injected with Ly6G antibody (1A8, 500μg) one day before and after wounding. Blood was extracted on wound day 2 and analyzed by flow cytometry for percent neutrophils (Gr1+, Ly6c+). *p = 0.0362 as calculated by two-way ANOVA. n.s., not significant. N = 2 vs 3 and 2 vs 2. Results are representative of at least two independent experiments. **j**. Neutrophil depletion from the wound bed is successful on small, but not large wounded mice. Mice wounded with small or large wounds (**h**.) were IP injected with Ly6G antibody (1A8, 500μg) one day before and after wounding. Wound beds were collected on wound day 2 and analyzed by flow cytometry for percent neutrophils (Gr1+, Ly6c+). Depletion was significant in the small wound, but not the large wound setting. *p = 0.0458 as calculated by two-way ANOVA. n.s., not significant. N = 2 vs 3 and 2 vs 2. Results are representative of at least two independent experiments.

Unexpectedly, on wound day 3, after receiving two IP injections of either Ly6G or Gr-1 antibodies, using our careful detection strategy, there was no apparent neutrophil depletion in the wound bed, as measured via flow cytometry (Fig. 3b). Depletion was also ineffective in other tissues after wounding, including the spleen and liver (Fig. S3). Looking one day earlier (wound day 2), neutrophil depletion was still ineffective in the wound bed, but there was a statistically significant depletion in the blood of the same mice (Fig. 3c). Although significant, the neutrophil depletion was less than 50%, far less efficient than the previous reports^52,54,55^. An important distinction between ours and previous work is that we employed depletion in the context of large wounds to fascia, prompting us to question whether large area wounding globally modifies neutrophil blood trafficking dynamics to result in depletion failure. Indeed, neutrophil depletion was very efficient in the blood of unwounded mice who had received an IP injection of the Ly6G (1A8 clone) antibody 24 hours earlier (Fig. 3d), but this depletion was abolished after wounding the mice (Fig. 3e). To explore this further, we repeated the injection scheme in Fig. 3a. With and without wounding mice, we examined the neutrophil depletion in the blood and wound bed at wound day 2. Consistent with our previous experiments, in the context of wounding, the Ly6G (1A8) neutralizing antibody achieved less than 50 percent neutrophil depletion in the blood. In contrast, mice that were not wounded, achieved substantially more efficient depletion in the blood (Fig. 3f). Unwounded skin does not typically contain neutrophils, so the Ly6G (1A8 clone) antibody injection had no effect (Fig. 3g). Wounding caused dramatic recruitment of neutrophils to the wound bed, as shown earlier (Fig 3b-c, e). Ly6G antibody injection could not overcome this to deplete neutrophils from the wound bed (Fig. 3g).

A previous report had success with antibody-mediated neutrophil depletion in a wounding model where small 2cm excisions were made^55^, leading us to question whether Ly6G (1A8 clone) antibody injection was more effective in smaller, full-thickness wounds. To address this, mice were injected with Ly6G (1A8 clone) antibody and then wounded with large wounds (1.25×1.25cm2), like the previous experiments, or smaller wounds (0.6×0.6cm2) (Fig. 3h). Neutrophil depletion efficacy was then assessed at wound day two in the blood and wound bed. Mice that had been given small wounds had significant neutrophil depletion in both the blood and wound bed, while those with the large wounds did not (Fig 3i-j). Taken together, these data suggest that antibody mediated neutrophil depletion, while efficient in certain non-wounding contexts, is not effective after the large full-thickness wounds necessary to induce WIHN—perhaps because of the unusually robust stimuli of neutrophil recruitment in this context.

### Mature Neutrophils inhibit wound induced hair neogenesis

Despite the minimal inhibition of neutrophils using conventional antibody depletion with large wounds, we sought to begin testing how neutrophils might modulate WIHN. We used the same antibody depletion scheme described (Fig 3a) but allowed the mice to recover for 21 days before quantifying WIHN via confocal scanning laser microscopy (CSLM) (Fig 4a)^56^. Antibody injection did not affect wound closure speed (Fig. S4). Despite minimal neutrophil depletion within the wound beds of mice post Ly6G (1A8 clone) injection (Fig 3b, c, e, g), the number of regenerated hair follicles was increased, suggesting that neutrophils may hinder WIHN (fold = 3.94) (Fig 4b).

**Figure 4:**
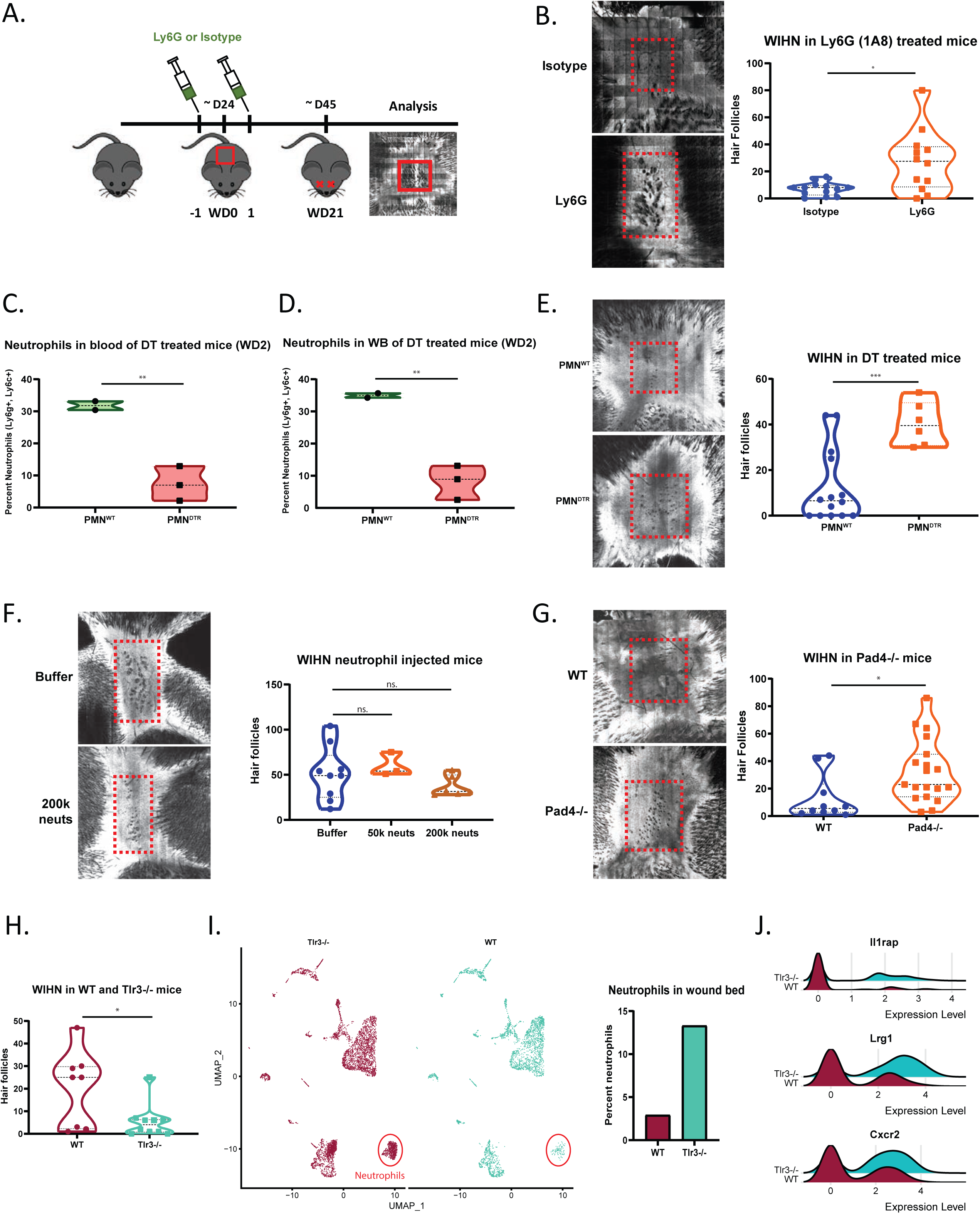
Mature neutrophils inhibit wound induced hair neogenesis. **a**. Schematic of neutrophil depletion via IP of Ly6G antibody (100μg) or isotype control. IP injections are done one day before and one day after wounding C57BL/J6 mice with 1.25×1.25cm^2^ square wounds. Wound induced hair neogenesis (WIHN) was measured 21 days after wounding via confocal scanning laser microscopy (CSLM). **b**. Neutrophil targeted Ly6G antibody injected mice exhibit increased WIHN (CSLM, images; fold = 3.94, p = 0.011, N = 10 vs 9). In each image, the dash red box indicates the area of hair follicle regeneration. **c**. Diphtheria toxin (DT, 250ng) injection in heterozygous ROSA26iDTR/MRP8-Cre-ires mice (PMN^DTR^ and PMN^WT^), following the injection scheme in **a**., successfully depletes neutrophils from the blood in mice with large wounds. Fold = −4.33. **p = 0.0098. N = 2 vs. 3. Results are representative of at least two independent experiments. **d**. Mice treated as in **c**. are depleted of neutrophils in their wound beds at wound day 2. Fold = −4.27. **p = 0.0069. N = 2 vs. 3. Results are representative of at least two independent experiments. **e**. PMN^DTR^ mice IP injected with diphtheria toxin (DT, 250ng) on wound days −1 and 1 exhibit increased WIHN (CSLM, images; fold = 3.23, ***p = 0.0010, N = 14 vs 6). In each image, the dash red box indicates the area of hair follicle regeneration. **f**. Regenerated hair follicles after the injection of 50k-200k purified neutrophils underneath the scab at WD7-8. (CSLM, images; n.s., not significant, N = 9 vs 3 vs 6). In each image, the dash red box indicates the area of hair follicle regeneration. **g**. Pad4^-/-^ mice defective in extracellular traps exhibit increased WIHN (CSLM, images; fold = 2.47, p = 0.026, N = 10 vs 19). In each image, the dash red box indicates the area of hair follicle regeneration. **h**. Tlr3^-/-^ mice exhibit decreased WIHN (fold = −3.75 p = 0.0218, N = 8 vs 10). **i**. The presence of increased neutrophils correlates with decreased WIHN in TLR3^-/-^ mice. scRNA-seq t-SNE plot shows differences between WT (red, 4150 cells) and Tlr3-/- (blue, 5648 cells) wound beds at wound day 10. The plots were generated via Seurat. The neutrophil cluster is circled in red. Percent of neutrophils are graphed to the right. **j**. Neutrophil associated gene expression is more pronounced within the neutrophils of Tlr3-/- mice, compared to WT. Generated in Seurat with RidgePlot function.

To surmount the difficulties with meager neutrophil depletion using Ly6G antibodies, we generated a transgenic mouse model for selective and inducible ablation of neutrophils upon injection of diphtheria toxin (DT)^57^. MRP8-Cre mice expressing Cre recombinase under the control of the neutrophil-associated human MRP8 promoter^58-60^ were crossed with ROSA-iDTR^KI^ mice, which have a Cre-inducible simian DT receptor (DTR)^61^. This generated mice with DTR expression restricted to mature neutrophils that suffer cell death selectively after the injection of DT. Two injections, one before wounding and one a day after, had no effect on control mice (PMN^WT^ mice: MRP8-Cre-, ROSA-iDTR^KI^) but substantially reduced neutrophil numbers in the blood (fold = −4.33) and wound beds (fold = −4.27) of PMN^DTR^ (MRP8-Cre+, ROSA-iDTR^KI^) mice at wound day 2 (Fig. 4c-d). We next followed the mice after neutrophil ablation in the early phase of wounding to find WIHN was substantially elevated (fold = 3.23) (Fig. 4e). The ablation of neutrophils later in the wound healing process yielded the same results (Wound days 6, 8, and 10) (Fig. S5). Antibody mediated depletion (1A8 clone) at these later time points had no effect on WIHN, suggesting that although effective in the blood, it is not able to target neutrophils already present within the wound bed (Fig. S6). After overcoming the technical hurdles above, these data suggest that neutrophils have a detrimental effect on the regeneration of hair follicles.

To further explore this possibility, we sought to try a gain-of-function approach, rather than the loss-of-function approach above. We injected purified neutrophils from the bone marrow of mice into the wound beds of mice during the re-epithelialization process. Although not significant, likely due to the high variability of WIHN, the addition of 200,000 neutrophils reduced the number of regenerated hair follicles (fold = −0.73) (Fig. 4f).

Finally, given the presence of NETs late in wound healing (Fig 2e), we sought to test the role of NETs in hair follicle neogenesis directly. We, therefore, tested WIHN in NET-deficient Pad4 KO mice, as employed in Figure 2e. In the absence of Pad4 and with dramatically reduced NETs, WIHN is enhanced (fold = 2.47, p = 0.026) (Fig. 4g). This suggests that NETs play a role in reducing the regenerative capacity of mice during the wound healing process. This correlates with the capacity of NETs to damage tissue in diseases such as small vessel vasculitis^62^, systemic lupus erythematosus^63-67^, rheumatoid arthritis^68^, and psoriasis^69^.

### Single-cell RNAseq correlation of mature neutrophils with poor WIHN

Double-stranded RNA sensing, mediated by Tlr3 and downstream effector pathways Il-6/ Stat3, has been shown to be critical for WIHN, with Tlr3-/- mice having substantially less regenerated hair follicles than their wild-type controls^34^. Given that TLR3 dsRNA sensing has been shown to be critical for neutrophil recruitment and NET production in a model for acute lung injury (ALI) and glomerulonephritis (GN), we wondered if Tlr3-/- mice paradoxically have increased neutrophil levels, contributing to lower WIHN^70,71^. As anticipated, Tlr3-/- mice had substantially less WIHN than their WT controls (fold = −3.75, p = 0.0218) (Fig. 4h, Fig. S7a). We performed single-cell RNA sequencing on wildtype and Tlr3-/- re-epithelialized wound beds of mice 10 days post wounding and prior to morphogenesis. Approximately 8014 sequenced cells met standard quality control metrics and were further analyzed in the Seurat R package^72^. Unsupervised clustering and UMAP non-linear dimensional reduction identified 18 cell clusters. Seurat generated conserved and differentially expressed genes to assign cluster identities. We found the mature neutrophil cluster to be substantially increased in the Tlr3-/- mice (Fig. 4i, Fig. S7b). This disparity is evident when looking at select differentially expressed neutrophil associated genes that are all elevated in Tlr3 −/- mice (Fig. 4j). Together, these data suggest that elevated mature neutrophil levels within Tlr3-/- mice correlate with their deficiency in WIHN.

## Discussion

The wound healing process is a careful balance of interconnected steps that must weigh the benefits of quick barrier repair, which leads to fibrous scarring, and more complete regeneration, that restores function and appearance. The role inflammation plays in regeneration and scarring is still being elucidated, but increasing evidence suggests that neither excess nor lack of inflammation supports regeneration. Fgf9-producing γ-δ T cells are critical for WIHN, infiltrating into wound bed immediately before re-epithelialization and onset of hair follicle regeneration^7^. Macrophages have also been shown to be important in the process, with their ablation eliminating WIHN^8-10^. Additionally, the injection of the dsRNA mimic poly(I:C), as early as 3 days after wound, dramatically enhances WIHN^34^. Intriguingly, spiny mice (*Acomys*) have a dramatically reduced inflammatory response post wounding, with less cytokines and virtually no macrophages until late in the process, when compared to laboratory mice (*Mus*), despite having substantially improved regeneration^73,74^. Consistent with this idea of a complex network of inflammatory cells influencing regeneration, we show that neutrophils persist in the wound bed after the acute inflammatory phase and—despite multiple correlations to the contrary-- have a detrimental effect on regeneration.

Using histological and flow cytometry techniques, we show that while abundant immediately after wounding, mature neutrophils remain in low levels within the wound bed after re-epithelization. These late stage neutrophils produce NETs, which are eliminated in mice that are deficient in Pad4, an enzyme critical for chromatin decondensation and NET formation^50,51^. Critically, Pad4-/- mice have increased WIHN, when compared with WT mice. Furthermore, antibody mediated and selective genetic neutrophil ablation dramatically boost WIHN. Single-cell RNA sequencing also revealed that mice deficient in dsRNA sensing Tlr3, who have severely reduced WIHN, have substantially more mature neutrophils present in the re-epithelized wound bed, immediately preceding regeneration. All these data suggest a model where neutrophils play an important role in defense against bacterial pathogens, but if they persist within the wound bed too long after barrier repair, regeneration is hindered. In the future, it will be interesting to see if selectively targeting NETs (e.g., PAD4 inhibitors, DNase I, N-acetylcysteine) enhance regenerative wound repair.

When attempting to deplete neutrophils to assess their effects on WIHN, we also discovered that the commonly used antibody depletion methods were not effective. Two neutrophil-specific antibodies, which recognize different epitopes of the GPI-linked differentiation antigen Ly6G, are commonly injected to rapidly deplete circulating neutrophils. The most widely used clone (1A8) specifically recognizes Ly6G, while the anti-granulocyte receptor-1 (Gr-1) antibody (RB6-8C5 clone) recognizes a heterodimer of Ly6G and Ly6C, making it partially target monocytes. Both antibodies efficiently depleted neutrophils in unwounded animals. When large full-thickness wounds (1.25×1.25cm^2^) were used, which are necessary to initiate WIHN, depletion levels in the blood were drastically reduced, while having negligible effects on neutrophils within the wound bed. Depletion efficacy was restored if smaller wounds were used (0.6×0.6cm^2^), albeit at reduced levels when compared to unwounded mice. These results suggest that wounding has a dramatic effect on recruiting neutrophil precursors for mobilization and release, overwhelming the standard ability of depletion antibodies to work effectively. This highlights the importance of rigorously testing antibody mediated depletion methods in mouse models, to ensure they are achieving appreciable depletion. Furthermore, it is important to carefully design neutrophil detection methods to assess depletion efficacy, particularly with flow cytometry. Robust depletion can be erroneously detected if similar Ly6G clones are used for both depletion and detection, due to epitope masking^53^. This can occur since identical monoclonal clones are sometimes given different clone numbers. Antibody manufacturers are increasingly manipulating antibodies to reduce undesirable features like non-specific binding, without affecting the epitope-binding variable region, yet renaming the clones, necessitating substantial effort to ensure non-overlapping epitope binding by both depletion and analytical flow antibodies.

Our work generates important areas of future investigation. One question is unraveling the paradox of why neutrophil infiltration signatures correlate with high WIHN, but mature neutrophils inhibit WIHN. One possible model is that a common upstream cue or factor both promotes WIHN and promotes neutrophil infiltration, but the latter serves to limit WIHN in favor of decreasing infection risk. Defining this common upstream signal will be important for future work. It will also be interesting to define the function of neutrophils within the re-epithelized wound since the barrier has been restored. Another important question is whether nuclear RNAs released in NETs have any function besides the general theorized one for released DNA. Though NETs inhibit WIHN, nuclear RNA release in NETs might have functions in separate physiological processes that will yield important insights. A final question is the identity of the neutrophil mobilization factor whose strength correlates to wound size. Potential signals include products of complement activation (C5a) or a number of small molecular weight C-X-C chemokines, which are recognized by Cxcr1 and Cxcr2^75,76^.

In summary, we here demonstrate a novel role for NETs and mature neutrophils to inhibit regeneration. Future studies will be important to further understand the biology of regeneration and test the capacity for neutrophil inhibition to promote regenerative healing.

## Acknowledgments

The authors also thank Conover Talbot Jr. (JHMI Deep Sequencing and Microarray Core) for assistance with microarray analysis.

## Methods

### Mouse Lines

All wild-type and control mice used for in vivo experiments were on the C57BL/6J background. All mice were age-matched and co-housed until 3-weeks of age. *Pad4* knockout mice were purchased from the Jackson Laboratory (B6.Cg-Padi4^tm1.1Kmow^/J, 030315). The diphtheria toxin (DT) mediated neutrophil ablative mice were generated by crossing ROSA26iDTR (C57BL/6-Gt(ROSA)26Sor^tm1(HBEGF)Awai^/J, 007900) and MRP8-Cre-ires/GFP (B6.Cg-Tg(S100A8-cre,-EGFP)1Ilw/J, 021614) from the Jackson Laboratory to get heterogeneous mice and genotyped according to their specifications. Mice who genotyped positive for Cre (MRP8-Cre+; ROSA-iDTR^KI^) were considered PMN^DTR^ mice, while those that were negative for Cre (MRP8-Cre-; ROSA-iDTR^KI^) were PMN^WT^ littermate controls^57^. Tlr3 knockout mice (B6N.129S1-Tlr3^tm1Flv^/J,009675) and C57BL/6NJ controls (005304) were purchased from the Jackson Laboratory. All mice were bred and housed at an American Association for the Accreditation of Laboratory Animal Care (AAALAC)-compliant facility, and all experimental procedures were reviewed and approved by the Johns Hopkins University Institutional Animal Care and Use Committee (IACUC).

### Wound Induced Hair Neogenesis (WIHN) Assay

All in vivo experimental surgical procedures were performed as previously characterized^7,16,34,37,77^. Briefly, after exposure to anesthesia (Baxter, Isoflurane), the dorsal side of 3-week-old (21 days) male and female mice were shaved. Surgical scissors were used to excise 1.25×1.25cm^2^ of skin, creating wounds deep into the fascia. Approximately 3 weeks after wounding (∼21 days), neogenic hair follicles in the re-epithelialized skin tissue were quantified using reflectance confocal scanning laser microscopy (CSLM) as previously published^34,37^.

### Neutrophil Depletion

C57BL/6J mice were depleted with 100 or 500μg of anti-Ly6G (Bio-X-Cell, 1A8 clone, BP0075-1) via intraperitoneal injection (IP) injection one day prior to and one day after wounding mice^52,54^. Select experiments also used 200μg of anti-Gr1 (Bio-X-Cell, RB6-8C5 clone, BP0075). LgG2a (Bio-X-Cell, BE0089) and IgG2b (Bio-X-Cell, BP0090) isotype controls were used for Ly6G or Gr1 experiments, respectively. DT depletion was done with PMN^DTR^ and PMN^WT^ littermate control mice that were IP injected with 250ng DT (Sigma-Aldrich). The injections were primarily done one day before and after wounding, or at wound days 6, 8, and 10.

### Flow Cytometry

Flow cytometry was used to access neutrophil depletion. Blood was collected via retro-orbital sinus bleeds, and red blood cells were lysed RBC lysis buffer (BioLegend, 420301). Wound beds were surgically removed, and cell suspensions were prepared by digesting the tissue in a cocktail consisting of Liberase TL (Roche, 5401020001) and DNase I (Sigma, DN25) in RPMI 1640 (Gibco, 11875093). Cells were washed and then Fc blocked (BioLegend, 101320), before staining with an antibody cocktail (Extended Table 1). Finally, cells were washed and resuspended in FACs buffer containing Propidium Iodide (Miltenyi, 130-093-233). All flow cytometry experiments were performed on a BD LSR II, and downstream analysis of data was performed using FlowJo.

### Neutrophil Extracellular Trap measurement

Wound beds were surgically removed at wound days 2 and 7, using a 6mm biopsy punch to remove excess tissue. Cell suspensions were prepared by digesting the tissue in a cocktail consisting of Liberase TL (Roche, 5401020001) and DNase I (Sigma, DN25) in RPMI 1640 (Gibco, 11875093). Cells were washed, and then Fc blocked (BioLegend, 101320), before staining with an antibody cocktail containing anti-MPO (Abcam, ab208670, 1:500) or respective isotype control (Abcam, ab172730, 1:500). Cells were then stained with a secondary Alexa Fluor 647 antibody (Abcam, ab150083, 1:2000). On the final wash, SYTOX green was added (ThermoFisher, S7020, 1:1000). This was performed on a BD LSR II, and downstream analysis of data was performed using FlowJo.

### 3’-end single-cell RNA-sequencing

The re-epithelialized wounds beds (wound day 10) of a Tlr3-/- and a C57BL/6NJ control mouse were excised, and cell suspensions were prepared by digesting the mouse skin tissue in a cocktail consisting of Liberase TL (Roche, 5401020001) and DNase I (Sigma, DN25) in RPMI 1640 (Gibco, 11875093). Propidium iodide and DAPI positive dead cells were removed via cell sorting with a BD FACSAria II. Single-cell libraries were prepared via a 10× Genomics Chromium Single-Cell Platform, followed by sequencing using Illumina NovaSeq 6000. The results were run through Cell Ranger pipeline software for sequence alignment and basic filtering. GEM generation, barcoding, cDNA amplification, library preparation, quality control, and sequencing were performed at the Genomics High Throughput Sequencing facility at Johns Hopkins School of Medicine.

Downstream analysis, after the Cell Ranger pipeline, was done using the Seurat R package. A standard pre-processing workflow was done, removing low quality cells or doublets, filtering unique feature count over 3750 and below 200, as well as filtering out cells with higher than 5% mitochondrial counts. This resulted in 4150 WT and 5648 Tlr3-/- cells for downstream bioinformatics. Expression matrices then underwent normalization, scaling, principal components analysis, and subsequent t-SNE analysis using Seurat packages. Seurat was then used to generate conserved genes, differentially expressed genes, feature plots, dot plots, and ridge plots. Cell clusters were then defined querying conserved genes and differentially expressed genes against the Immgen gene expression database (www.immgen.org) using the interactive tool “My Gene Set.”

### Histology

Biopsies from mouse skin tissue were removed and fixed in 4% paraformaldehyde overnight and then transferred to 70% ethanol. Samples were then submitted to the Johns Hopkins Oncology Tissue Services Core facility where they were embedded in paraffin. Tissue sections were obtained at 4μm thickness and mounted onto glass slides, followed by hematoxylin and eosin (H&E) staining.

### Immunofluorescence and immunohistochemistry

Immunofluorescence microscopy was performed on de-paraffinized tissue sections that received heat-induced antigen retrieval using Target Retrieval Solution (Agilent Dako, S169984-2). After washing and permeabilization in TBS-T universal buffer (0.2% Triton X-100 in tris-buffered saline), sections were blocked at room temperature in 5% goat, donkey, or fetal bovine serum with 1% bovine serum albumin. Tissue sections were then incubated overnight at 4°C with primary antibodies (Extended Table 2) in Antibody Diluent (Agilent Dako, S080983-2). Following a wash step, sections were incubated in fluorescent-dye conjugated secondary antibodies diluted in antibody diluent for 1 hour at room temperature. After final washing, sections were mounted with VECTASHIELD® Hardset™ Antifade Mounting Medium with DAPI (Vector Laboratories, H-1500) for nuclear staining. All imaging was done on a DFC365FX (Leica) at 20x and 40x magnifications.

### U1 in situ hybridization

U1 in situ probes were designed and ordered in the Stellaris Probe Designer (Biosearch Technologies) (Extended Table 3). Tissue sections were de-parafinized and stained following Biosearch Technologies Stellaris RNA FISH protocol for Formalin-Fixed Paraffin-Embedded Tissue. Briefly, tissue sections were washed in Wash Buffer A (Biosearch Technologies, SMF-WA1-60), before adding 200μl hybridization buffer (Biosearch Technologies, SMF-HB1-10) containing the U1 probe and covering the tissue with a glass coverslip. The slides were then incubated overnight in a humid box at 37^°^C. Slides were then immersed in Wash Buffer A in the dark at 37^°^C for 30 minutes, allowing the coverslips to float off. Slides were then washed for 5 minutes with Wash Buffer B (Biosearch Technologies, SMF-WB1-20), before sections were mounted with VECTASHIELD® Hardset™ Antifade Mounting Medium with DAPI (Vector Laboratories, H-1500) for nuclear staining. All imaging was done on a DFC365FX (Leica) at 63x magnifications.

### Microarray, RNA-seq and proteomic analysis

Proteins from the wound center and wound edges were analyzed by proteomics, as previously described^37^. Briefly, we used Orbitrap Fusion Tribrid mass spectrometer (Thermo Scientific) for protein profiling and obtained the protein expression level by the MS Amanda algorithm. Standard bioinformatics procedures were performed, including standardization of gene expression, the definition of differentially expressed genes, and GO enrichment analysis. For both Rnasel-/- and WT mice, total RNA was isolated from mouse tissue at the time of scab detachment from the wound (∼10 days post-wounding). RNA was submitted to the JHMI Deep Sequencing & Microarray core facility and profiled using the Affymetrix Clariom™ S mouse array platform, according to the manufacturer’s protocols. Gene chips were scanned, generating CEL pixel intensity files, which were processed and analyzed using Partek^®^ Genomics Suite™ software, and the Robust Multichip Analysis (RMA) algorithm was used for normalization. For SPF and GF analysis, total RNA from early wound bed skin (∼WD12) was submitted to the JHMI Transcriptomics and Deep Sequencing Core. The 1.0ST exon sequencing of mouse RNA was performed according to the manufacturer’s standard protocol. The raw affymetrix CEL data was standardized using Robust Multichip Analysis (RMA) algorithm for comparison.

### Quantification and Statistical Analysis

All in vivo and in vitro experiments were performed in at least individual instances. Univariate statistical analysis was performed using Student’s t-test, and multivariate analysis was performed using ANOVA. All statistical analyses and graphical representations were generated using GraphPad Prism software. Statistical significance is defined as p-values <0.05 derived from the standard error of mean calculations.

**Extended Table 1.**

| Name | Host | Fluorophore | Manufacturer/Product # |
| --- | --- | --- | --- |
| MHCII (IA/IE) | Rat | BV421/Pacific Blue | BioLegend/107631 |
| CD3 | Rat | BV510/AmCyan | BD/740147 |
| Ly6C | Rat | FITC | BD/553104 |
| Ly6G | Rat | PE | BioLegend/127607 |
| CD45 | Rat | PE-Cy5.5 | Invitrogen/35-0451-82 |
| CD115 | Rat | PE-Vio770 | BioLegend/135523 |
| CD11c | Hamster | APC | BioLegend/117310 |
| CD11b | Rat | APC-Cy7 | BD/557657 |

**Extended Table 2.**

| Name | Host | Dilution | Company/ Product # |
| --- | --- | --- | --- |
| MPO | Goat | 1:200 | Abcam/ab208670 |
| F4/80 | Rat | 1:200 | Abcam/ab6640 |
| Ly6G | Rat | 1:200 | BioXCell/BP0075-1 |
| H3Cit | Rabbit | 1:500 | Abcam/ab5103 |
| Alexa Fluor® 488<br>Anti-Goat IgG (H+L) | Rabbit | 1:1000 | Invitrogen/A27012 |
| Alexa Fluor® 488<br>Anti-Rabbit Ig (H+L) | Goat | 1:1000 | Invitrogen/A-11008 |
| Alexa Fluor® 594<br>Anti-Rabbit IgG (H+L) | Goat | 1:1000 | Invitrogen/A-11037 |
| Alexa Fluor® 488<br>Anti-Rat Ig (H+L) | Goat | 1:1000 | Invitrogen/A-11006 |
| Alexa Fluor® 594<br>Anti-Rat Ig (H+L) | Donkey | 1:1000 | Invitrogen/ A-21209 |

**Extended Table 3.**
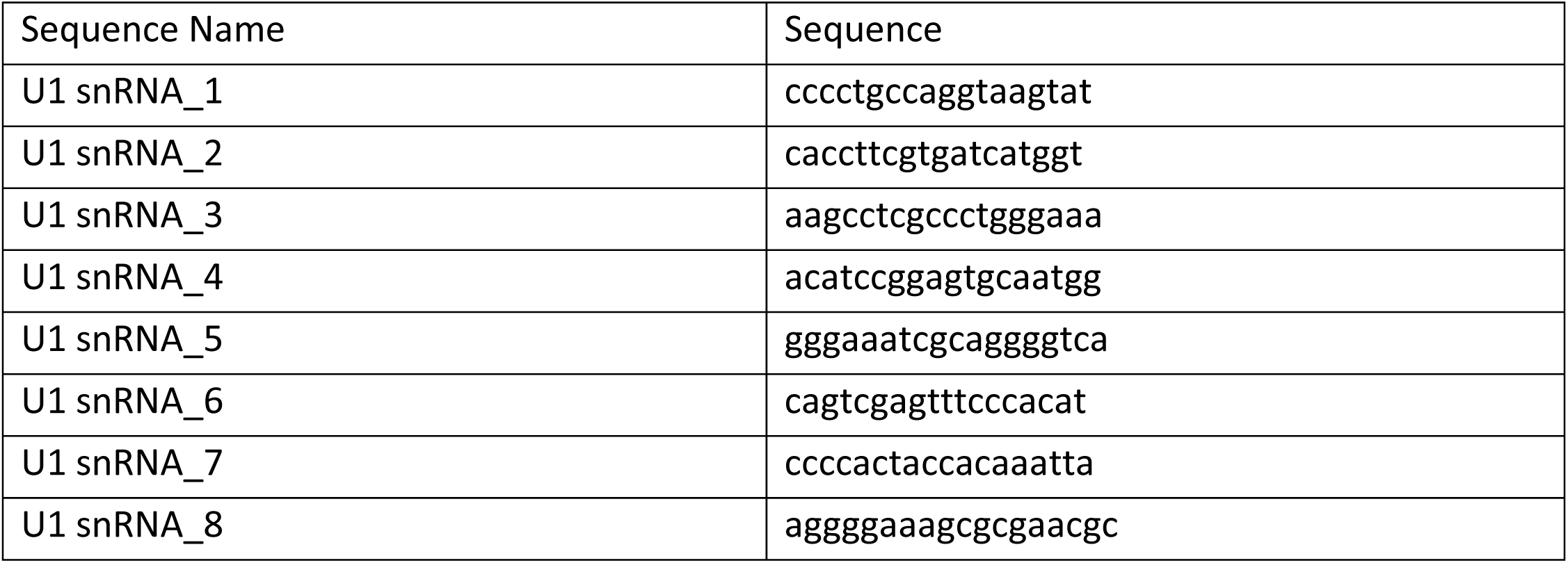

## Supplementary Figure Legends

**S1:** Macrophage (F4/80) levels are largely absent from the blood and low in the wound bed during the early phase of healing, but increase dramatically at wound day 11, as measured by FACS. ***p < 0.004, as calculated by two-way ANOVA. n.s., not significant. N = 2 vs 4. Results are representative of at least two independent experiments.

**S2:** Ly6G 1A8 antibody clone (Bio-X-Cell) masks the Ly6G epitope, preventing detection by the Ly6G REA526 clone (Miltenyi, engineered 1A8 clone) and partially preventing detection by the Gr1 RB6-8C5 clone (Bio-X-Cell) antibodies. Blood was extracted from a single C57BL/6J and stained sequentially with combinations of two antibodies, to test Ly6G epitope masking, before detecting by flow cytometry. If only stained with the Ly6G (REA526) or Gr1 (RB6-8C5), the neutrophil population is detected normally (13.7% or 13.2%, respectively). If you first stain with Ly6G (1A8 clone), followed by Ly6G (REA526) or Gr1 (RB6-8C5), you get varying degrees of Ly6G antibody masking. Being a derivative of the 1A8 clone, the REA526 Ly6G binding site is almost completely blocked by the prior incubation with the Ly6G 1A8 clone (1.10%), making the pair unusable for neutrophil depletion experiments. Prior incubation with the Ly6G 1A8 clone followed by Gr1 (recognized an epitope of Ly6G/Ly6C) shifts the neutrophil population to the left, but they remain distinct from the negatively stained cells (12.7%). With careful gating, this makes depletion with the Ly6G (1A8 clone) antibody and detection with the Gr1 (RB6-8C5) antibody possible. The neutrophil population is boxed in red.

**S3:** Antibody mediated Neutrophil depletion is not efficient in the blood, spleen, liver, or wound bed, two days after wounding. The percent neutrophils (CD11b+, Ly6c+), at wound day 2, present in the blood, spleen, liver, and wound beds of mice IP injected with Ly6G antibody (500μ, 1A8) one day before and one day after given 1.25×1.25cm^2^ full thickness wounds. The neutrophil depletion is statistically insignificant, as calculated by two-way ANOVA. n.s., not significant. N = 4 vs 4, except for the Blood and Wound bed samples, which were 7 vs. 7.

**S4:** The wound closure rate was not affected by neutrophil depletion via IP of Ly6G antibody (100μg) or isotype control. IP injections were done one day before and one day after wounding C57BL/J6 mice with 1.25×1.25cm^2^ square wounds. N = 4 vs 4. Results are representative of three independent experiments.

**S5**. PMN^DTR^ mice IP injected with diphtheria toxin (DT, 250ng) on wound days 6, 8, and 10 exhibit increased WIHN (CSLM, images; fold = 3.28, **p = 0.0085, N = 13 vs 5).

**S6: a**. Schematic of neutrophil depletion via IP of Ly6G antibody (500μg) or isotype control. IP injections are done on wound day 8 and 10, after wounding C57BL/J6 mice with 1.25×1.25cm^2^ square wounds. Wound induced hair neogenesis (WIHN) was measured 21 days after wounding via confocal scanning laser microscopy (CSLM). **b**. Neutrophil depletion in the blood is successful on wound day 9, one day after antibody injection (Ly6G, 1A8), via FACs. The neutrophil depletion is statistically significant. ****p < 0.0001, as calculated by two-tailed Student’s t-test. Fold = −87.9. N = 2 vs 3. **c**. Late stage Ly6G antibody injected mice exhibit normal WIHN (CSLM, images; n.s., not significant, N = 3 vs 3). In each image, the dash red box indicates the area of hair follicle regeneration.

**S7: a**. Tlr3^-/-^ mice exhibit decreased WIHN (representative CSLM images, N = 8 vs. 10). In each image, the dash red box indicates the area of hair follicle regeneration. **b**. Neutrophil associated genes used to identify the neutrophil cluster in UMAP non-linear dimensional reduction, via Seurat R package, of WT and Tlr3-/- mice.

